# Bovine colostrum-derived extracellular vesicles impair cancer cell proliferation through transcriptional repression

**DOI:** 10.1101/2025.10.07.680845

**Authors:** CB Huesa-Carballo, B Puga, S Bravo, B Hermida, I Raña, MT Antelo, R López-López, M Abal, J Barbazán

## Abstract

Milk-derived extracellular vesicles (EVs) are a promising source of molecules with therapeutic potential. Bovine colostrum is particularly enriched in EVs, which carry a unique cargo of proteins involved in immune regulation, development, and cellular signaling. However, despite promising bioactive effects in various fields, little is known about their potential as anti-cancer agents. Here, we demonstrate that colostrum-derived EVs (Col-EVs) exert a potent and specific anti-proliferative effect on gastrointestinal cancer cell lines, independent of apoptosis induction. Using a multi-modal approach combining proteomics and functional assays, we show that Col-EVs induce a reversible proliferative arrest through transcriptional repression and chromatin and nuclear remodeling. Col-EV treatment leads to widespread dysregulation of RNA processing and transcriptional machinery, including the downregulation of splicing factors and chromatin regulators essential for cell cycle progression. These molecular changes are accompanied by chromatin compaction, nuclear reorganization, and cytoskeletal remodeling. Notably, these effects do not induce cell death and are reversible upon EV removal, suggesting a modulatory mechanism rather than cytotoxicity. Furthermore, Col-EVs enhance the efficacy of DNA-targeting chemotherapies such as 5-fluorouracil, indicating their potential as adjunctive agents in cancer treatment. Overall, our findings reveal that Col-EVs can selectively and reversibly suppress cancer cell proliferation by reprogramming transcriptional and nuclear architecture, offering a natural, biocompatible strategy for modulating tumor growth and sensitizing cancer cells to conventional therapies.

## Introduction

Milk is a nutrient-rich biological fluid that plays a central role in neonatal development, immune system maturation, and gut homeostasis (1–3). Beyond its classical nutritional components, which include proteins, fats, carbohydrates, vitamins, and minerals, milk contains extracellular vesicles (EVs), nano-sized lipid bilayer particles capable of transferring bioactive cargos such as proteins, RNAs, lipids, and metabolites between different cells, which don’t have self-replicative abilities (4,5). These milk-derived EVs (mEVs) have been implicated in a wide range of physiological processes, including modulation of gut barrier integrity (6,7), immune responses (2,8,9), and microbiota composition (3,5,10), thereby shaping health outcomes from infancy into adulthood.

Recent studies have demonstrated that mEVs are uniquely resistant to gastrointestinal degradation (6), enabling their oral bioavailability and functional uptake by gut epithelial cells and even systemic distribution to distal organs, including the liver and tumors. This stability, coupled with their biocompatibility and low immunogenicity, positions mEVs as promising candidates for oral nanotherapeutics and drug delivery vehicles, with applications ranging from inflammatory bowel disease and gut-liver axis disorders to systemic modulation of immune responses and cellular signaling (11–13). While the potential of mEVs in supporting gut health and reducing inflammation is increasingly recognized, their potential application as anti-cancer agents remains underexplored. Interestingly, some studies have shown that bovine milk-derived EVs can reduce primary tumor burden in colorectal and breast cancer models by inducing tumor cell senescence (14).

Bovine colostrum, the first milk produced post-parturition, is particularly enriched in EVs containing high levels of immune-modulatory proteins, antimicrobial peptides, and growth-regulatory factors (5,15). These colostrum-derived EVs (Col-EVs) encapsulate a unique protein and RNA cargo that reflects the biological function of colostrum as a system supporting early immune training, epithelial barrier maturation, and tissue development. Indeed, this specialized composition suggests that Col-EVs may possess distinct biological activities compared to mature milk-derived EVs (16,17), with potential for differentially influencing cancer cell proliferation and survival. However, the mechanistic understanding of if and how Col-EVs could impact cancer cell proliferation remains limited.

In this study, we investigate the anti-proliferative potential of bovine Col-EVs on gastrointestinal cancer cells. We hypothesize that Col-EVs induce a reversible growth arrest in cancer cells by interfering with transcriptional programs and promoting chromatin compaction, thereby suppressing proliferation without inducing apoptosis. Using a combination of proteomics, advanced imaging, and functional assays, we dissect the mechanistic pathways involved in this phenotype, providing insights into the therapeutic potential of Col-EVs as natural, orally available bio-nano modulators in cancer therapy.

## Results

### Comprehensive Characterization of bovine Col-EVs

In order to understand the protein composition of EVs from bovine colostrum (Col-EVs), we adapted an EV isolation protocol (18) based on sequential ultracentrifugation combined with different filtration steps (Supplementary Fig. 1). Nanoparticle tracking analysis (NTA) revealed a size distribution with a major peak around 100-150 nm and a long tail extending to 400 nm, consistent with the expected profile for exosome-enriched EV populations (Fig. 1A, left panel). This heterogeneity likely reflects both exosomal and small microvesicle populations. Quantification of replicate samples showed consistent particle concentration and size distribution (Fig. 1A, center and right), indicating the robustness of the isolation protocol. Moreover, Col-EVs were visualized using transmission electron microscopy (TEM), confirming their vesicular morphology. Vesicles appeared spherical and sized in the 100–200 nm range (Fig. 1B), supporting their classification as small EVs. Of note, other components such as lipid or cassein aggregates were detected as part of the sample, as expected due to their high abundance in this sample type (Supplementary Fig. 2A). To further profile surface markers, we employed Single-Particle Interferometric Reflectance Imaging Sensor (SP-IRIS) analysis using antibody-based capture and detection. Using capture antibodies against CD63, CD81, and CD9, we quantified the expression of canonical tetraspanins on individual vesicles (Fig. 1C). CD9 was the most abundantly expressed tetraspanin, present on the majority of particles. CD81 and CD63 were detected at lower levels. As well, we found that CD63 and CD81 positive vesicles co-expressed at different degrees other tetraspanins, while the vast majority of CD9-isolated vesicles did not co-expressed other markers (Fig. 1D). These results validate the isolation methodology and indicate phenotypic heterogeneity within the isolated Col-EV population.

**Figure 1.**
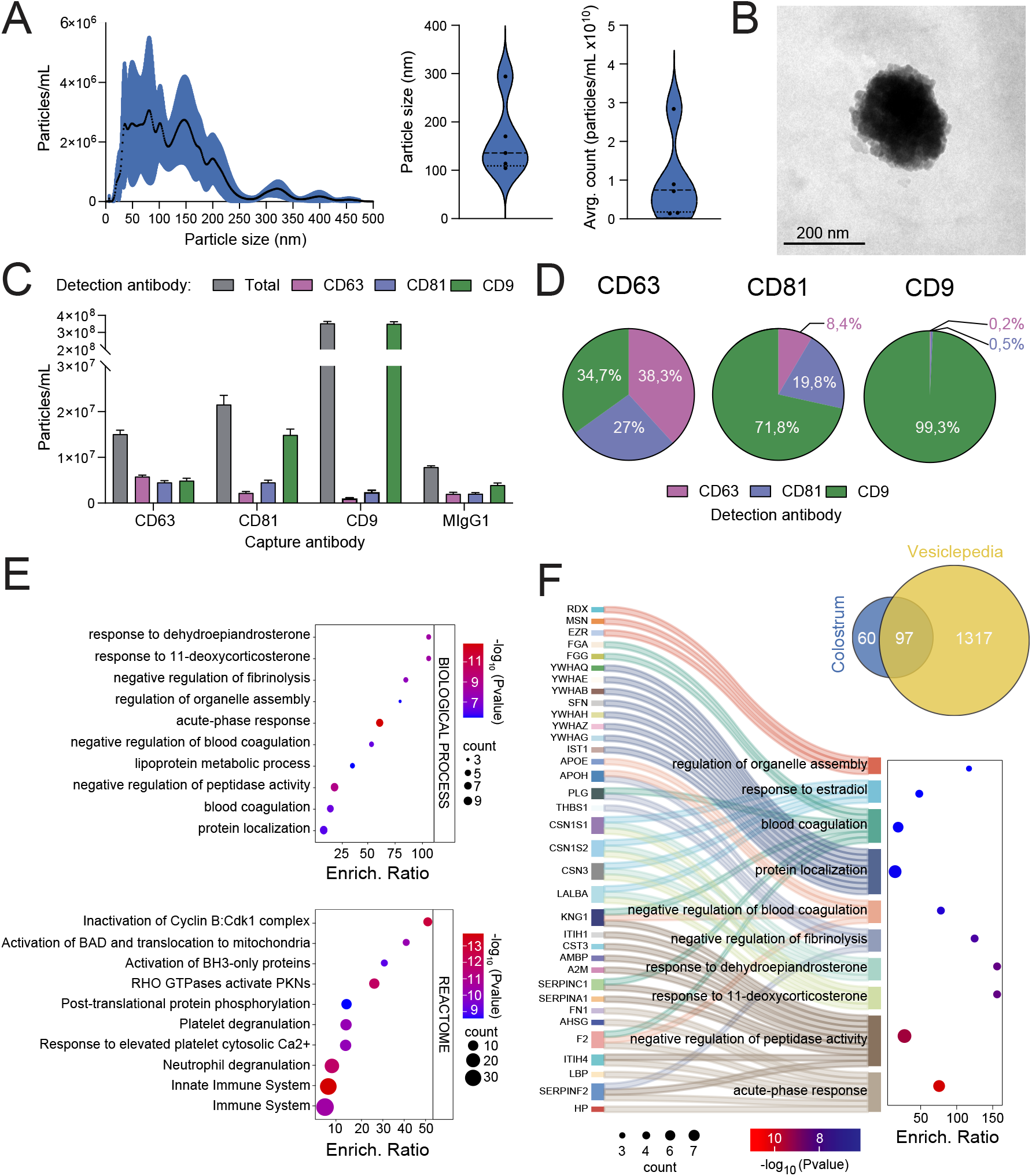
Characterization of bovine colostrum-derived EVs. (A) Left panel: nanoparticle tracking analysis (NTA) showing the size distribution Col-EVs, with a primary peak around 100–150 nm. (N=6) Middle and right panels: violin plots of the quantification of particle size and concentration (N=5), respectively. (B) Representative transmission electron microscopy (TEM) images confirming the spherical morphology and expected size range (100–200 nm) of isolated Col-EVs. (C) Quantification of particle concentration using SP-IRIS (N=3). MIgG1 indicates the signal obtained using an isotype control. Error bars: SD. (D) Co-expression analysis showing heterogeneity in tetraspanin profiles across the Col-EV population (N=3). (E): Gene Ontology (Biological Process) (upper panel) and Reactome pathway enrichment analyses (lower panel) of the Col-EV DDA proteomic analysis. (F) Venn diagram and Sankey analysis demonstrating overlap with Vesiclepedia and highlighting functional categories of identified proteins. For EV proteomics samples were run in triplicates.

To further characterize the proteome of Col-EVs, we performed proteomic analyses using data-depending acquisition (DDA) mass spectrometry. Using this approach, over 150 specific proteins were detected with less than 1% of FDR (False Discovery Rate). Further analyses showed an enrichment in pathways associated with inflammation, coagulation and complement activation, as revealed by both Gene Ontology (Biological Process) and Reactome pathway analyses (Fig. 1E, Supplementary Excel File). These results are in line with previous reports of colostrum proteomics, which consistently described colostrum as a fluid that contains high levels of antimicrobial peptides, immunoglobulins, complement proteins, and factors involved in wound healing and epithelial maturation (15–17) Notably, several proteins involved in organelle assembly and steroid responses were also enriched, indicating broader functional roles for these EVs. The detection of proteins related to organelle assembly and steroid responses suggests that Col-EVs may also participate in intracellular signaling and metabolic priming, potentially supporting early tissue development and endocrine regulation. Comparison with the Vesiclepedia database (19) revealed a high percentage of proteins associated with EVs (Fig. 1F, Venn diagram and Supplementary Excel File), reinforcing the presence of EVs in the sample. Such proteins included fibrinogen subunits (FGA, FGB, FGG), coagulation factors, and stress-response proteins such as HP, APOE, and LBP. Sankey visualization linked these proteins to functional categories such as regulation of organelle assembly, response to hormones (e.g., estradiol), and negative regulation of proteolytic activity. (Fig. 1F and Supplementary Excel File)

Overall, these data indicate that bovine Col-EVs possess a specialized protein cargo with potential immunomodulatory, structural, and regulatory properties.

### Col-EVs inhibit gastrointestinal cancer cell proliferation without inducing cell death

To determine the potential biological effects of Col-EVs on cancer cells, we treated a panel of gastrointestinal epithelial cancer cell lines and primary cancer-associated fibroblasts (CAFs) with Col-EVs. Alamar Blue viability assays performed 72 hours post-treatment revealed a significant decrease in cell number/metabolic activity in colorectal (LoVo) and pancreatic (PANC1) cancer cells, while patient-derived CAFs were unaffected by Col-EVs, suggesting selective targeting of epithelial cancer cells without elevated toxicity to the surrounding stromal compartment. As well, and to assess the impact of EVs on cell growth dynamics, we performed time-lapse microscopy assays over a period of 72 hours. Col-EV-treated PANC1 and LoVo cells exhibited a marked inhibition of proliferation, as they failed to expand into a monolayer during time-lapse imaging (Fig. 2B and Supplementary Video 1). Quantitative analysis of the cancer cell area filled over time confirmed a drastic reduction in growth compared to controls (Fig. 2C). In addition to a reduction in proliferation, we observed that Col-EVs induced a distinct morphological phenotype, forming compact, spheroid-like aggregates which were particularly evident in LoVo cells (Fig. 2B), suggesting a decreased cell-substrate adhesion and increased cell-cell compaction. To better visualize these structural changes, we performed confocal imaging of F-actin and DAPI-stained LoVo cells. While in control conditions cells displayed a flat, epithelial monolayer morphology, Col-EV treated cells exhibited a rounded appearance and vertical multilayering, as revealed by orthogonal sectioning of the confocal images (Fig. 2D). This phenotype is consistent with a dewetting-like response and reflects the loss of adhesion and polarity associated with EV-induced growth arrest.

**Figure 2.**
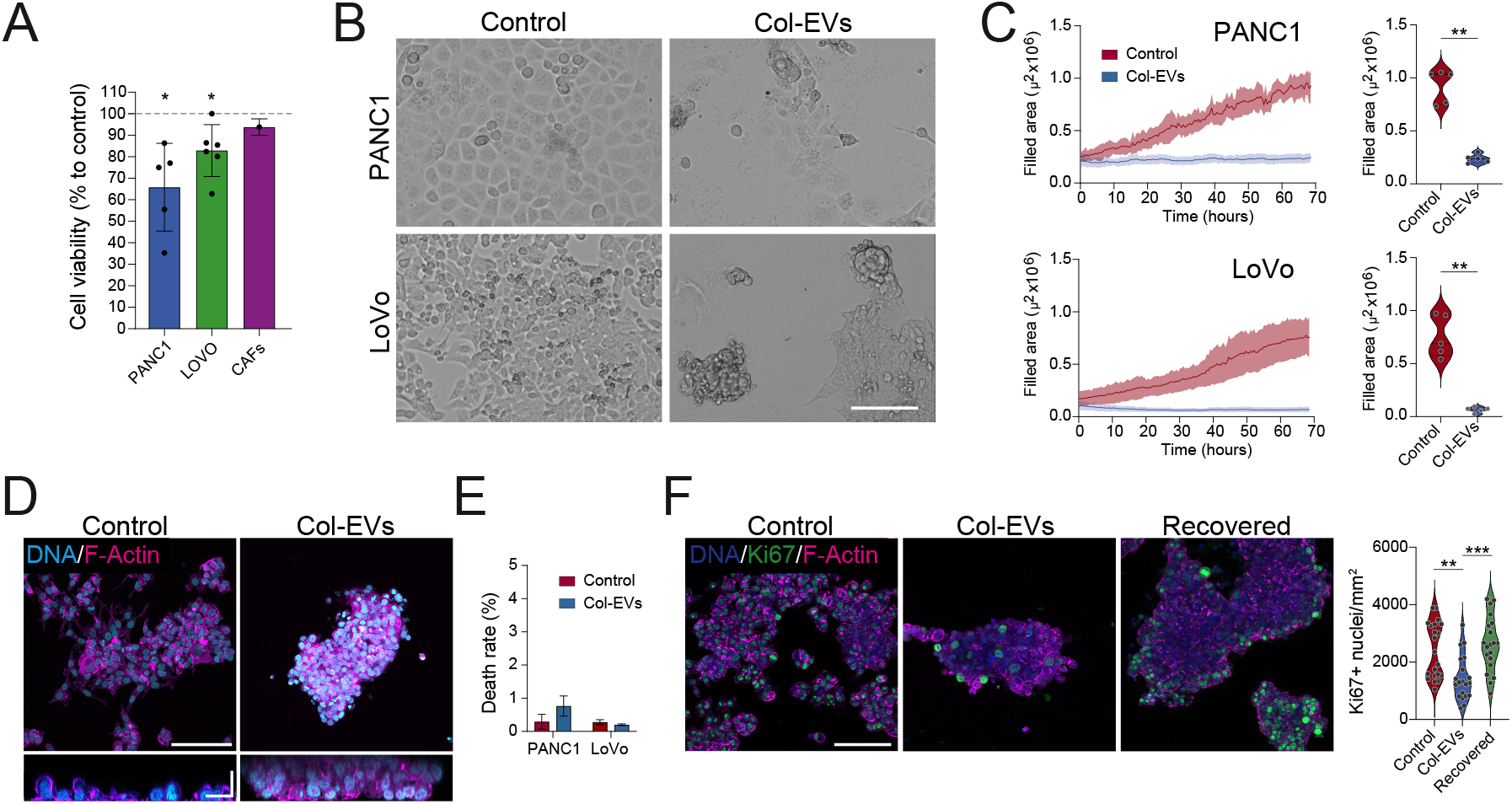
Col-EVs selectively impair cancer cell proliferation without inducing cell death. (A) Viability assays in colorectal (LoVo) and pancreatic (PANC1) cancer cells, and in patient-derived CAFs upon Col-EV treatment. Error bars: SD. *p<0.05 (N=5 for PANC1, N=6 for Lovo, N=1 for CAFs). (B) Representative time-lapse images of PANC1 and LoVo cells 72h after either control or Col-EV treatment. (C) Left panels: quantification of PANC1 and LoVo filled area over time from videomicroscopy experiments. Scale bar: 100 µm. Right panels: violin plots of the quantification of total filled area at t=72h post-treatment. **p<0.01. Data from n=5 independent positions/condition for N=1 experiment. (D) Representative confocal images of F-actin (magenta) and DAPI-stained (cyan) LoVo cells demonstrating vertical multilayering and reduced spreading upon Col-EV treatment. Scale bar: 50 µm for main images and 20 µm for orthogonal sections. (E) Cell death analysis of PANC1 and LoVo cells treated or not with Col-EVs. Error bars: SD. (N=2). (F) Left panel: representative images of Ki67-stained (green) control, Col-EV treated and recovered LoVo cells. F-actin (phalloidin): magenta. DNA (DAPI): blue. Scale bar: 50 µm. Right panel: quantification of Ki67 positive cells per area (mm^2^). **p<0.01; ***p<0.001. Data from n=20 images from N=2 independent experiments.

To determine whether the observed effects were due solely to growth arrest or also involved cell death, we performed cell death stainings, which revealed no significant differences between control and Col-EV treated cells (Fig. 2E), indicating that the treatment did not induce apoptosis. Contrarily, Ki67 immunostaining revealed a substantial reduction in the number of actively cycling cells under EV treatment (Fig. 2F). Notably, when EVs were removed from the culture medium, Ki67 positivity and monolayer organization were largely restored (Fig. 2F), demonstrating that the anti-proliferative effects of Col-EVs are reversible.

Overall, these results indicate that colostrum-derived EVs selectively suppress the proliferation of epithelial cancer cells without inducing cell death, instead promoting a reversible growth-arrested state. This is accompanied by striking changes in cellular architecture and adhesion, suggesting that Col-EVs interfere with both proliferative and mechanical cues in cancer cells.

### The anti-proliferative effects of Col-EVs are specific and linked to transcriptional dysregulation programs

To ensure that the observed biological effects were attributable specifically to EVs and not to other non-vesicular protein content present in the ultracentrifugation pellet, we designed a control experiment comparing the biological activity of Col-EVs and a protein-rich fraction. For this, we repeated the cell viability assays using Col-EVs isolated in the 100,000×g (100k) fraction and compared their effects to those of material recovered in the previous 70,000×g (70k) intermediate pellet from the same sample. Viability assays showed that only the 100k fraction significantly reduced proliferation in both PANC1 and LoVo cells (Fig. 3A), indicating that the anti-proliferative activity is specific to the Col-EV population and not attributable to contaminating proteins or lipids present in the bulk pellet. Of note, the 70k fraction also contains soluble proteins such as casein and lipids, as well as vesicles (Supplementary Fig. 2B). which could differ in their protein cargo due to the differential effects observed over cancer cells. Indeed, proteomic analyses of both fractions revealed a total of 28 proteins unique to the 100k Col-EV sample. These proteins include nuclear proteins such as histones H3-3A and H3-5, transcriptional regulators CDK1 and CDK13, and annexins, among others (Fig. 3B and Supplementary Excel File). Pathway enrichment analyses revealed cell cycle regulation, FOXO-mediated transcription, and G2/M checkpoints as some of the main functions in which proteins specific to Col-EVs were found to be involved, which could be linked to the observed effects over cell proliferation (Fig. 3C and Supplementary Excel File).

**Figure 3.**
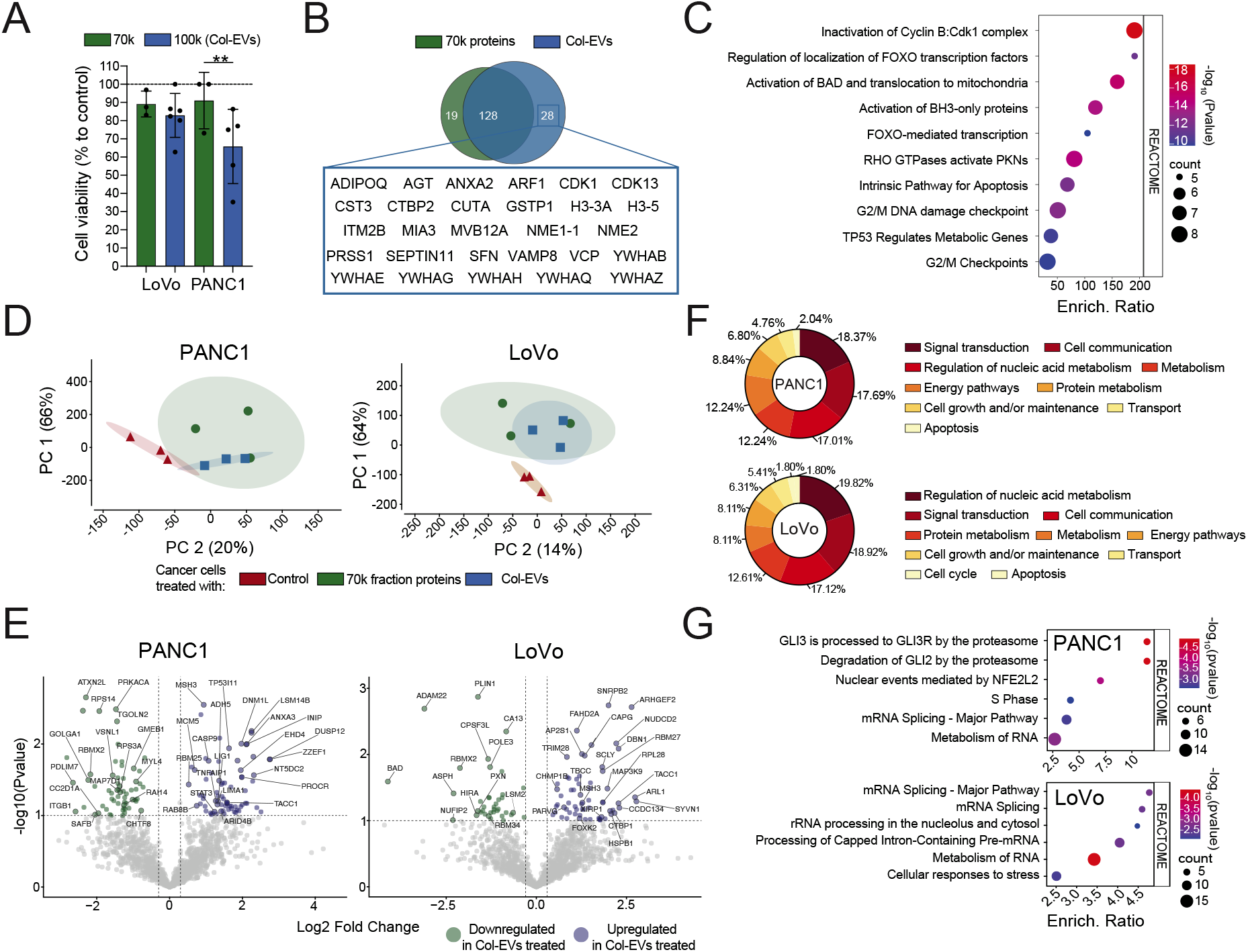
Col-EVs-specific proteins induce growth arrest in cancer cells through a mechanism compatible with transcriptional repression. (A) Viability assays comparing the effects of 70k (protein-rich) and 100k (Col-EV-enriched) fractions on LoVo and PANC1 cells. Error bars: SD. **p<0.01. Data shows at least 3 independent experiments per condition (B) Venn diagram displaying unique proteins identified in the Col-EV fraction. (C) Reactome pathway enrichment of Col-EV exclusive proteins showing involvement in cell cycle regulation, FOXO-mediated transcription, and G2/M checkpoint control. For EV proteomics samples were run in triplicates. (D) SWATH-MS proteomics principal component analysis (PCA) of control, 70k and Col-EV treated PANC1 and LoVo cells. Data obtained from 3 independent experiments per condition. (E) Volcano plots showing widespread dysregulation of proteins in PANC1 and LoVo cells upon treatment with Col-EVs. Fold change levels were calculated using 70k-treated cells as controls. Thresholds: P value: 0.01; Fold Change: 1.3/-1.3. Data obtained from 3 independent experiments. (F–G) Functional enrichment analyses (F:Funrich; G: Reactome) highlighting the top 6 dysregulated pathways in PANC1 and LoVo cells upon Col-EV treatment, compared with the treatment with the 70k protein fraction. Data obtained from 3 independent experiments.

To better understand the specific effects of Col-EVs on cancer cells, we performed quantitative whole-cell SWATH-MS proteomics on LoVo and PANC1 cells treated with either the 70k fraction or Col-EVs. Principal component analysis (PCA) showed that, as expected, the major differences were observed between control cells and those treated with both fractions. However, samples treated with the 70k fraction and Col-EVs also clustered separately from each other in both cell lines (Fig. 3D), indicating that Col-EVs exert a distinct and specific effect on cancer cells beyond that of the protein-rich 70k pellet. Volcano plot analyses revealed that treatment with Col-EVs led to a widespread dysregulation of protein expression across both LoVo and PANC1 cells (Fig. 3E and Supplementary Excel File). Col-EV treatment altered the abundance of key regulators of gene expression and cellular architecture. This included downregulation of RNA splicing factors (e.g., RBM25, SNRPB2), chromatin-associated enzymes (e.g., DNMT1, PRMT5), and structural proteins involved in mitotic progression (e.g., TACC1). In parallel, we observed dysregulation of proteins involved in cytoskeletal dynamics and adhesion, including CAPG, a modulator of actin filament capping; MYH9, a non-muscle myosin heavy chain important for cell contractility; and integrin β1 (ITGB1), a major mediator of cell-ECM interactions, consistent with the observed cellular phenotypes.

To further explore the biological relevance of these proteomic shifts, we performed pathway enrichment analysis using FunRich and Reactome. In both LoVo and PANC1 cells, treatment with Col-EVs led to a significant enrichment of dysregulated proteins involved in transcriptional regulation, RNA metabolism, and cell cycle progression (Fig. 3F and Supplementary Excel File). Notably, Reactome analysis highlighted the disruption of critical pathways such as mRNA splicing, S phase progression, and the processing of capped pre-mRNAs (Fig. 3G and Supplementary Excel File), processes that are essential for the transcriptional and post-transcriptional control of gene expression.

Collectively, these results suggest that Col-EVs mediate their anti-proliferative effects by targeting core transcriptional and RNA processing pathways, thereby depriving cancer cells of the molecular machinery required to sustain cell cycle progression.

### Col-EVs induce reversible chromatin compaction and nuclear reorganization

To investigate whether the observed transcriptional repression could be linked to structural nuclear changes, we examined chromatin organization using high-resolution confocal microscopy. Upon treatment with Col-EVs, LoVo cells displayed marked alterations in nuclear architecture, including prominent DAPI-bright foci. Images quantification confirmed a reduction of the DNA-occupied nuclear area as well as an increase in the DAPI signal standard deviation per nucleus, both indicative of a chromatin condensation phenotype (Fig. 4A, B). These changes were largely absent in control cells, in which interphase nuclei showed smooth and homogeneous DNA distribution. Concordantly with the observed Ki67 positivity ratios (Fig. 2F), the nuclear DNA distrubution returned to normal levels upon Col-EV withdrawal (Fig. 4A, B), reinforcing the transitory/reversible effect of Col-EVs.

**Figure 4.**
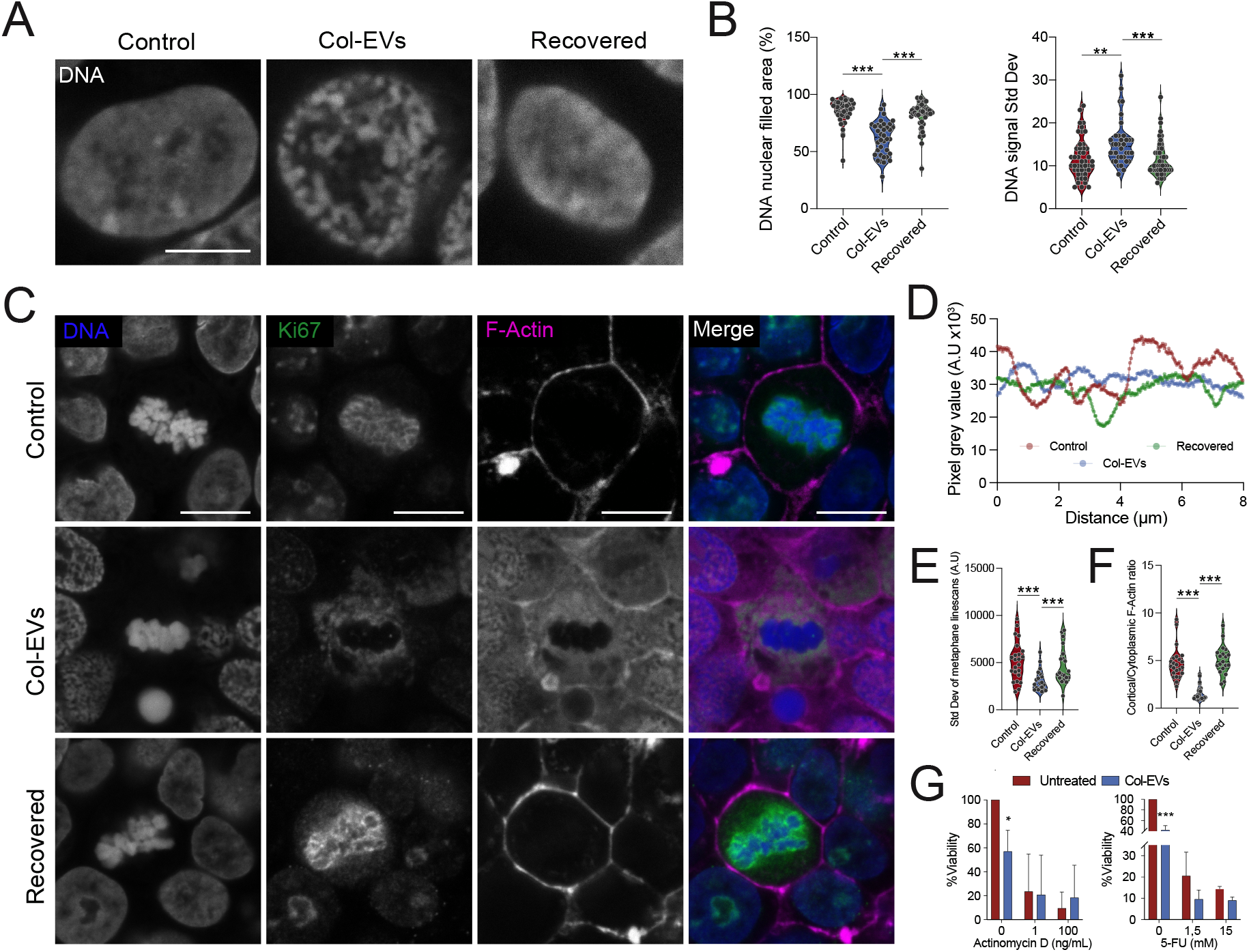
Col-EVs induce reversible chromatin compaction and nuclear remodeling. (A) Representative confocal images of DAPI-stained nuclei from LoVo cells, control, Col-EV treated or recovered. Scale bar: 5 µm (B) Quantification of the % nuclear area covered by DNA (left panel), as well as the standard deviation of the nuclear signal/nucleous, confirming reversible chromatin compaction upon EV removal. Data represents at least 35 analyzed nuclei from n=2 independent experiments. **p<0.01, ***p<0.001; Kruskal-Wallis ANOVA test (C) Representative confocal images of mitotic figures in LoVo cells, stained for DNA (DAPI, blue), Ki67 (green) and F-Actin (Phalloidin, magenta), from control, Col-EV treated or recovered conditions. Scale bar: 10 µm (D) Representative line-scan analysis of DAPI signal intensity in metaphasic chromosomal plates from images in panel. (E) Quantification of the standard deviation of DAPI intensity across line scans in metaphase plates, where lower standard deviation in Col-EV-treated cells indicates reduced peak variability consistent with hypercompaction and loss of individualized chromosome structure. (F) Quantification of F-Actin cortical/cytoplasmic ratios. For E and F data represents at least 20 analyzed nuclei from n=2 independent experiments. ***p<0.001; Kruskal-Wallis ANOVA test. (G) Viability assays in PANC1 cells treated with increased doses of actinomycin D (left panel) or 5-FU (right panel), with or without Col-EVs. (n=at least 4 independent replicates). Error bars: SD. *p<0.05; ***p<0.001, ANOVA test.

Despite the drastic changes in chromatin organization observed, we could observed cells undergoing mitosis in the Col-EV treated condition, evidenced by the presence of metaphasic chromosomal plates (Fig. 4C). However, when closely analysing those mitotic figures after performing line-scan analysis of DAPI signal intensity across nuclei, we observed that Col-EV treated cells showed an aberrant chromosomal hypercompaction, with no individual chromosomes being distinguished (Fig. 4C-E). This effect could be a result of the constitutive presence of heterochromatin upon Col-EV treatment, which does not prevent cells from undergoing M-phase, but likely resulting in aberrant incomplete division that leads to proliferation arrest. Indeed, Ki67, which normally locate around individual chromosomes acting as a surfactant to prevent chromosome clumping and aberrant division (20), was found to be located around the main metaphasic chromosome mass (Fig. 4C and Supplementary Fig. 3), likely unable to properly localize due to chromatin compaction missregulation, and a readout of inefficient abnormal cell division. All these changes were accompanied by a drastic reorganization of F-actin pools, not able to properly localize to the cell cortex during division and remaining misslocalized over the entire cytoplasm in Col-EV treated cells (Fig. 4C, F). As expected, cells reverted to the initial phenotype upon Col-EV retrieval, supporting the reversibility of the phenotype.

Finally, to assess whether the phenotype induced by Col-EVs was primarily driven by transcriptional arrest, we treated cells with increasing doses of actinomycin D, a potent inhibitor of RNA synthesis, alone or in combination with Col-EVs, and measured cancer cell viability. As expected, actinomycin D alone caused a marked reduction in proliferation. However, co-treatment with Col-EVs did not further enhance this effect (Fig. 4G, left panel), suggesting that global transcriptional inhibition by actinomycin D may render cells unresponsive to additional transcriptional interference by Col-EVs. Notably, the anti-proliferative effect of Col-EVs alone was less pronounced than that of actinomycin D, supporting the idea that Col-EVs exert a milder, more modulatory effect on transcription, possibly accounting for the reversibility of the phenotype observed upon EV removal. In contrast, when cells were co-treated with Col-EVs and 5-fluorouracil (5-FU), a chemotherapeutic agent whose primary mechanism of action involves inhibition of thymidylate synthase and incorporation into DNA, we observed a significant enhancement of cytotoxicity compared to 5-FU alone (Fig. 4G, right panel). This suggests that while Col-EVs primarily act through transcriptional and post-transcriptional dysregulation, their combination with DNA-targeting agents like 5-FU may produce a synergistic effect on cell proliferation.

In summary, Col-EVs induce reversible chromatin compaction and nuclear disorganization, which could lead to defective mitosis and impaired proliferation. These structural and transcriptional alterations contribute to growth arrest and might enhance sensitivity to DNA-targeting therapies like 5-FU.

## Discussion

Our study demonstrates that bovine colostrum-derived extracellular vesicles (Col-EVs) possess potent anti-proliferative effects on gastrointestinal cancer cells, mediated through transcriptional repression and chromatin remodeling. These effects are specific to epithelial tumor cells, reversible upon EV withdrawal, and not associated with classical apoptosis or overt cytotoxicity. This selective and transient inhibition of proliferation suggests a regulatory, rather than destructive, interaction between Col-EVs and recipient cancer cells.

The phenotypic changes observed in PANC1 and LoVo cells, namely reduced proliferation, chromatin compaction, nuclear reorganization, and cytoskeletal disassembly, point to a global reprogramming of cellular architecture and gene expression. Proteomic analysis confirmed that Col-EV treatment leads to the dysregulation of RNA processing and transcription-related proteins, many of which are essential for cell cycle progression and metabolic maintenance. Notably, the presence of histones in Col-EVs raises the intriguing possibility that these proteins could be transferred into recipient cancer cells and incorporated into chromatin, promoting a repressive epigenetic landscape. This may underlie the observed chromatin condensation and transcriptional silencing, and aligns with previous reports of horizontal transfer via EVs (21–23). However, we cannot exclude the possibility that part of the observed phenotype is also mediated by the transfer of genetic material. EVs are known to carry nucleic acids, including double-stranded DNA fragments, nucleosome-associated DNA, and regulatory small RNAs such as miRNAs and lncRNAs, and these cargo molecules could influence recipient cells through multiple mechanisms.

One of the most striking features of the col-EV-induced phenotype is its reversibility. Upon removal of EVs from the culture medium, cancer cells restored normal morphology, Ki67 expression and proliferation as well as DNA architecture. This reversible quiescent-like state bears resemblance to tumor dormancy or transient growth arrest, phenomena with therapeutic implications for controlling residual disease and preventing relapse. The capacity to induce a pause in proliferation without causing cell death could be particularly beneficial in reducing tumor burden while sparing surrounding tissues from cytotoxic damage. Additionally, we demonstrate that pharmacological inhibition of transcription using actinomycin D phenocopies the effects of Col-EVs, while co-treatment with 5-fluorouracil (5-FU) reveals heightened sensitivity in EV-treated cells. These findings suggest that Col-EVs may sensitize cancer cells to DNA-damaging or anti-metabolic drugs by interfering with transcriptional and chromatin regulatory programs. From a mechanistic standpoint, this convergence on transcriptional repression underscores a key vulnerability in cancer cells that could be exploited therapeutically.

The structural changes induced by Col-EVs extend beyond the nucleus. We observed significant disorganization of the actin cytoskeleton, including reduced cortical F-actin and altered cell spreading. These changes may impair mechanical signaling, polarity, and motility, factors that are crucial in metastasis and invasion. Although our study did not directly assess migration or invasion, the morphological shift toward a less adherent, more compact phenotype suggests potential effects on epithelial-to-mesenchymal transition (EMT) and metastatic competence. Interestingly, previous reports have shown that, while milk-derived EVs prevent primary tumor growth, they could be linked to an acceleration of tumor progression and metastasis (14). In light of our results, such observations could differ in the case of colostrum-derived EVs, guaranteeing further exploration.

From a translational perspective, Col-EVs offer several unique advantages. They are naturally occurring, biocompatible, and derived from a widely consumed food product. Their ability to survive gastrointestinal transit and exert systemic effects supports the feasibility of oral or local delivery. Furthermore, the selective targeting of epithelial tumor cells without impacting patient-derived fibroblasts, suggests a favorable safety profile. The identification of distinct EV cargo and downstream proteomic signatures also provides opportunities for biomarker discovery.

Future studies should focus on in vivo validation of these findings, including biodistribution, therapeutic efficacy, and safety in immunocompetent models of gastrointestinal cancer. It will also be critical to define the key molecular mediators within the EV cargo responsible for transcriptional repression and chromatin remodeling. Targeted engineering or enrichment of these components could enhance the therapeutic utility of Col-EVs and lead to next-generation EV-based biotherapeutics.

In summary, our findings uncover a novel mechanism by which bovine colostrum-derived EVs modulate cancer cell biology. By inducing a reversible, transcriptionally repressed state characterized by chromatin condensation and cytoskeletal remodeling, Col-EVs represent a promising, naturally derived strategy for controlling tumor proliferation. This work expands the functional landscape of dietary EVs and provides a strong foundation for their development as therapeutic agents in oncology.

## Materials and Methods

### Isolation of Colostrum-Derived Extracellular Vesicles (Col-EVs)

For Col-EVs isolation, first, bovine colostrum (40 mL) was diluted 1:1 with sterile phosphate-buffered saline (PBS) and centrifuged at 3,000×*g* for 15 minutes at 4 °C to remove fat and cellular debris. The upper fat layer was carefully removed, and the pellet was discarded. The resulting supernatant was transferred to ULTRACLEAR™ ultracentrifuge tubes (Beckman Coulter) and subjected to sequential differential ultracentrifugation using a 32SWT swinging bucket rotor (Beckman Coulter) as follows:

- 12,000×*g*, 1h, 4°C (discard fat layer)
- 30,000×*g*, 1h, 4°C
- 70,000×*g* (first spin), 1h, 4°C
- 70,000×*g* (second spin), 1h, 4°C

The final supernatant was filtered through 0.45 μm and 0.22 μm pore size syringe filters (Millipore). A final ultracentrifugation step was then performed at 100,000 × *g* for 2h at 4°C. The resulting was resuspended in 2.5 mL of sterile, serum-free DMEM/F12 medium, filtered through a 0.22 μm syringe filter and stored at –80 °C until further use.

### Col-EVs characterization

To characterize Col-EVs, the following techniques were used:

#### Nanoparticle Tracking Analysis (NTA)

Isolated Col-EVs were diluted 1:20 in sterile filtered PBS in order to reach the optimal concentration range for particle detection. Measurements were performed using a nanoparticle tracking analysis system (NanoSight). All measurements were conducted at room temperature (25.0 °C) under constant flow conditions with a flow rate of 40 μL/min. Measurement time was approximately 60 seconds per capture. A Blue488 laser and an sCMOS camera were used for all samples.

Acquisition and analysis parameters including camera level, shutter speed, gain, frame rate (FPS), number of frames and detection threshold, were individually adjusted for each sample to optimize particle visualization and tracking. The viscosity was set to that of water (0.9 cP). Data was analyzed using the manufacturer’s software (NTA 3.4.003), with automatic settings for blur size and maximum jump distance.

#### Single-Particle Interferometric Reflectance Imaging Sensor (SP-IRIS)

In order to evaluate the expression of EV-specific surface markers we used the SP-IRIS (ExoView) technology. For this, sample dilutions were prepared based on the concentration of particles/ml obtained previously through NTA, with the optimal concentration, as recommended by the manufacturer, being 10^9^ particles per chip for incubations. Each kit comprises 8 chips (pre-coated with capture antibodies or customizable), various solutions, and detection antibodies anti-tetraspanins, specifically anti-CD9 (Alexa Fluor® 488), anti-CD81 (Alexa Fluor® 594), and anti-CD63 (Alexa Fluor® 647).

For data analysis, fluorescent cut-offs were modified from 200 a.u. (for red, far-red and green channels) and from 400 a.u. (for blue channel) to limit the number of detected particles on MIgG to 50 events (for red, far-red, and green) or 100 events (for blue) following the technical instructions of the manufacturer using the ExoView Analyzer v3.2 (Unchained Laboratories).

#### Transmission Electron Microscopy (TEM)

Samples were prepared on carbon-coated copper grids for transmission electron microscopy analysis. Briefly, Col-EV samples were diluted 1:10 in ultrapure water, followed by gentle vortexing to ensure homogeneity. Using fine forceps, a carbon-coated copper grid was carefully placed on a Parafilm. Then, 10 µL of the diluted sample was applied onto the grid surface. Excess liquid was gently wicked away using filter paper to avoid damaging the grid or sample. The grids were then left to air dry at room temperature. To enhance contrast, grids were subsequently stained with one drop of 1% phosphotungstic acid solution for 1 minute, followed by one drop of ultrapure water to wash off excess stain. The grids were then allowed to dry completely before imaging on a transmission electron microscope (model JEM2010) at the microscopy unit of the University of Santiago de Compostela (CACTUS).

### Cell viability assays

Cell lines (LoVo and PANC1) were maintained in complete culture media under standard incubation conditions. LoVo cells were cultured in DMEM/F12 supplemented with 10% fetal bovine serum (FBS) and 5 mL penicillin/streptomycin (P/S), while PANC1 cells were maintained in high-glucose DMEM with 10% FBS and 5 mL P/S.

For viability assays, cells were seeded one day prior to treatment at densities of 10,000 cells per well in 96-well plates and 50,000 cells per well in 24-well plates. On day 1, culture medium was removed and replaced with a mixture of Col-EVs and fresh medium. For 96-well plates, 50 µL of Col-EVs suspension was diluted in 150 µL of medium; for 24-well plates, 167 µL of Col-EVs suspension was diluted in 500 µL of medium. Cells were incubated with treatments for 96 hours. Proliferation/viability was then assessed using alamarBlue cell viability reagent (ThermoFisher Scientific). Cells were incubated for 3 h with a 1:10 alamarBlue solution (diluted in complete medium), and then fluorescence was measured in a spectrophotometer using 580/590 nm (excitation/emission) filter settings.

Controls included untreated cells, cells treated with bovine serum albumin (BSA) and an additional centrifugation fraction, to assess for the specificity of the effect of Col-EVs.

### Live-cell imaging

Cells were seeded into 24-well plates one day prior to imaging and incubated overnight at 37 °C with 5% CO_2_. Time-lapse live-cell imaging was performed using a Celldiscoverer 7 automated microscope (Zeiss) equipped with a temperature- and CO_2_-controlled incubation chamber (37 °C, 5% CO_2_) to maintain physiological conditions throughout the experiment. Imaging was carried out using a Plan-Apochromat 5×/0.35 objective lens. Brightfield images were acquired every 30 minutes over a 69-hour period, yielding a total of 138 time points per position. For each well, five distinct positions were imaged. Brightfield contrast was used with a light source intensity set at 10%.

### EV treatment and flow cytometry analysis

LoVo and PANC1 cells were seeded at a density of 5 × 10^4^ cells per well in 24-well plates one day prior to treatment. Cells were then treated with Col-EVs for 96 hours under standard incubation conditions (37 °C, 5% CO_2_). After incubation, cells were harvested by trypsinization (0.05% trypsin-EDTA, 5 min, 37 °C), collected by centrifugation, and fixed in 4% paraformaldehyde (PFA) for 20 minutes at room temperature. Fixed cells were stained with Zombie NIR™ Fixable Viability Dye (BioLegend) at a 1:1000 dilution in PBS for 30 minutes at room temperature in the dark.

Flow cytometry acquisition was performed on a FACSCanto™ III cytometer (BD Biosciences) using BD FACSDiva™ Software Version 9.0. The APC laser was used for Zombie NIR detection, with voltage settings of 297 for APC, 187 for FSC-A, and 313 for SSC-A. A total of 10,000 events were recorded per sample. Data was analyzed using standard gating strategies with FlowJo software (v10.7.1, BD Life Sciences).

### Proteomic analysis

Proteomic analysis was carried out at the proteomics core facility of the Health Research Institute of Santiago de Compostela. For this, LoVo and PANC1 cells were seeded in 6-well plates (10^5^ cells per well) and then treated for 48 hours with either control, col-EVs, and 70k fraction extracts. Treatments were conducted in triplicates.

Proteomic analyses were performed as previously described (24,25) Briefly, for EV proteomics, 15 μg of Col-EVs or 70k fraction proteins were mixed with a lysis buffer in ratio 1:1 boiled and concentrated in a resolving 10% SDS-PAGE gel (26). For cell line proteomics, after treatment, cells were lysed and protein quantified and preconcentrated in a SDS-PAGE gel, followed by protein band staining (Sypro Ruby fluorescent staining; Lonza, Basel, Switzerland) and band excision. Gel pieces were reduced (10 mM DTT; Sigma-Aldrich, St. Louis, MO, USA) and alkylated (55 mM iodoacetamide; Sigma-Aldrich, St. Louis, MO, USA). Then, we performed in-gel tryptic digestion, as previously described (24,25). Peptides were extracted [50% ACN/0.1% TFA (×3) and ACN (×1)], pooled, concentrated in a SpeedVac, and stored at _−_20 °C.

#### Mass spectrometric analysis (DDA acquisition)

DDA analysis was made as previously described (24).Briefly, digested peptides (over 4µg of each sample: Col-EVs, 70K fraction, LoVo, PANC1) were separated using Reverse Phase Chromatography. A 90 minutes gradient ranging from 2% to 90% mobile phase B, was created using a micro liquid chromatography system (Eksigent Technologies nanoLC 400, Sciex) coupled to a high-speed Triple TOF 6600 mass spectrometer (Sciex) with a micro flow source. Data acquisition was performed by a TripleTOF 6600 System (Sciex, Foster City, CA) using a data-dependent analysis (DDA) workflow. Source and interface conditions were the following: ionspray voltage floating (ISVF) 5500 V, curtain gas (CUR) 25, collision energy (CE), 10 and ion source gas 1 (GS1) 25. Instrument was operated with Analyst TF 1.7.1 software (Sciex, USA). Switching criteria was set to ions greater than mass to charge ratio (m/z) 350 and smaller than m/z 1400 with charge state of 2–5, mass tolerance of 250 ppm and an abundance threshold of more than 200 counts per second (cps). Previous target precursor ions were excluded for 15 s. The instrument was automatically calibrated every 4 hours using tryptic peptides from PepCalMix as external calibrant.

After MS/MS analysis (MS2 data), data files were processed using ProteinPilot™ 5.0.1 software from Sciex, which uses the algorithm Paragon™ for database search and Progroup™ for data grouping. Data was searched using a Bovine specific Uniprot database (https://www.uniprot.org/) specifying iodoacetamide at cysteine alkylation as variable modification and methionine oxidation as fixed modification. False discovery rate was performed using a non-lineal fitting method, displaying only those results that reported a 1% Global false discovery rate or better. (27)

#### Protein quantification by SWATH (Sequential Window Acquisition of all Theoretical Mass Spectra)

For label-free quantitative proteomics, we performed an MS analysis by sequential window acquisition of all theoretical mass spectra (SWATH-MS), as previously described (24,25). First, a unique peptide pool was created by mixing equal amounts of peptides from each sample type (LoVo, PANC1). This pool was analysed by LC-MS/MS on a TripleTOF® 6600 LC-MS/MS system via a data-dependent acquisition (DDA) method in order to create a SWATH-MS spectral library. Only proteins and peptides with <1% FDR were included in this library (27). Then, 4 µg of peptides derived from the individual samples were analyzed by SWATH-MS method. SWATH-MS acquisition was performed on a TripleTOF® 6600 LC-MS/MS system via a data-independent acquisition (DIA) method. The whole 400 to 1250 m/z range was covered in 100 steps with spectral windows of variable width (1 m/z overlap). Peaks extraction was carried out with PeakView software (version 2.2; Sciex, Redwood City, CA, USA) and scored using the PeakView SWATH Acquistion MicroApp (version 2.0; Sciex, Redwood City, CA, USA). The integrated peak areas were exported to the MarkerView software (version 1.3, Sciex, Redwood City, CA, USA). To ensure a more accurate comparison between samples, well-known endogenous peptides were used during data alignment to compensate for small variations in both mass and retention times. The amount of each protein in every sample was calculated as the averaged area sums of 10 peptides per protein and 7 transitions per peptide. Then, an averaged MS peak area of each protein was calculated. Data normalisation was carried out with the most likely ratio normalisation (MLR) method. As part of the initial analysis, the MarkerView software (version 1.3, Sciex, Redwood City, CA, USA) also allowed a principal component analysis (PCA) to see how well each protein distinguishes between groups. We considered the proteins differentially expressed whit a *p*-value *<*0.01 and Fold change from 1.3/-1.3.

The mass spectrometry proteomics data have been deposited to the ProteomeXchange Consortium via the PRIDE (28) partner repository with the dataset identifier PXD066002 for Col_EVs and 70k fraction, and PXD066009 for treated cell lines.

Functional analyses were performed using the FunRich open access software (Functional Enrichment analysis tool) for functional enrichment and interaction network analysis (http://funrich.org/index.html) (29) and Gene Ontology analyses were performed using Genecodis4 (https://genecodis.genyo.es/) (30).

Volcano plots were generated using VolcaNoseR (31) and bubble plots were built using SRplot (31).

### Immunofluorescence staining and imaging

LoVo and PANC1 cells were seeded at a density of 5 × 10^4^ cells per well in 24-well plates one day prior to treatment. Cells were then treated with Col-EVs for 96 hours under standard conditions (37 °C, 5% CO_2_). Bovine serum albumin (BSA) was used as a vehicle control. Following treatment, cells were fixed with 4% paraformaldehyde (PFA) for 20 minutes at room temperature and subsequently washed with PBS.

An additional recovery condition was included in which cells were treated with Col-EVs for 96 hours, followed by a 48-hour incubation period in complete medium (medium change performed to remove col-EVs), and subsequently processed for staining as described.

#### Staining

Fixed cells were stained with DAPI (1:200) (ThermoFisher), phalloidin-CruzFluor™ 633 (1:500) (Santa Cruz Biotechnology Ref. sc363796), and Ki-67 (D3B5) Rabbit mAb (Alexa Fluor^®^ 488 Conjugate) (1:200) (Cell Signaling Technology Ref. #11882) for 1 hour at room temperature in the dark, followed by three washes with PBS.

#### Confocal Imaging

Stained conditions were imaged using an inverted confocal microscope (LEICA SP8) using laser lines 405, 488 and 633 nm and the following objectives: HC PL APO CS2 40x/1.30 OIL and HC PL APO CS2 63×/1.4 OIL. Images were processed using ImageJ (v2.14.0/1.54f).

### Image analysis

Image analysis was performed using Fiji/ImageJ (v2.14.0/1.54f). Brightfield images were converted to 8-bit format, and contrast was enhanced using the “Enhance Contrast” function with 0.35% saturated pixels and the *Normalize* option enabled, applied uniformly across all images. The background was subtracted using a rolling ball radius of 35 pixels. Binary masks were generated using the “Make Binary” function with Otsu thresholding, assuming a dark background. Particle analysis was conducted using the “Analyze Particles” tool, and the following parameters were extracted and summarized for each image: count, area, average size, percent area, mean intensity, mode, perimeter, and integrated density (IntDen).

### Comparative drug response assay

PANC1 cells were seeded in 96-well plates at a density of 10,000 cells per well and allowed to adhere overnight. Cells were then treated under co-treatment conditions for 96 hours, with all conditions performed in triplicate.

The following drug concentrations were tested:

- Actinomycin D: 1 ng/mL and 100 ng/mL
- 5-Fluorouracil (5-FU): 1.5 mM and 15 mM

After 96 hours of treatment, cell proliferation and viability were assessed using the alamarBlue™ Cell Viability Reagent (Thermo Fisher Scientific). Cells were incubated with a 1:10 dilution of alamarBlue in complete medium for 3 hours. Fluorescence was then measured using a microplate reader at 580 nm excitation and 590 nm emission.

### Data Analysis

Data visualization and statistical analyses were performed using GraphPad Prism 10. Statistical tests were selected based on data distribution. Experimental designs as well as the number of biological and technical replicates are specified in the corresponding figure legends. A p-value < 0.05 was considered statistically significant.

## Acknowledgements

We thank María Pardo and Nerea Lago (SP-IRIS/ExoView platform, IDIS Santiago), for their valuable assistance with EV characterization. We thank Nerea Lago for her reading and insightful feedback on the manuscript. Jorge Barbazán acknowledges funding from the Asociación Española Contra el Cáncer (AECC) through the *INVES246505BARB* contract. This work was funded with projects from the Spanish Ministry of Science, Innovation and Universities through the *Proyectos de Generación de Conocimiento* program (PID2023-152440OA-I00 for Jorge Barbazán, and PID2023-150296OB-I00 to Miguel Abal)

