## Supplementary Information for "Bovine colostrum-derived extracellular vesicles impair cancer cell proliferation through transcriptional repression"

### Supplementary Figures and figure legends

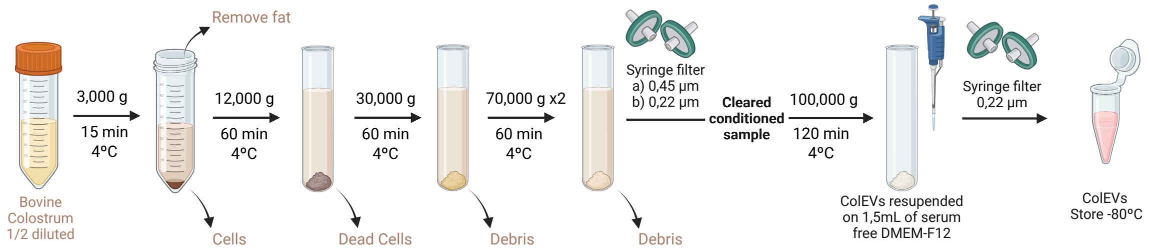


**Supplementary Figure 1.** Schematic representation of the EV isolation process from samples of bovine colostrum.

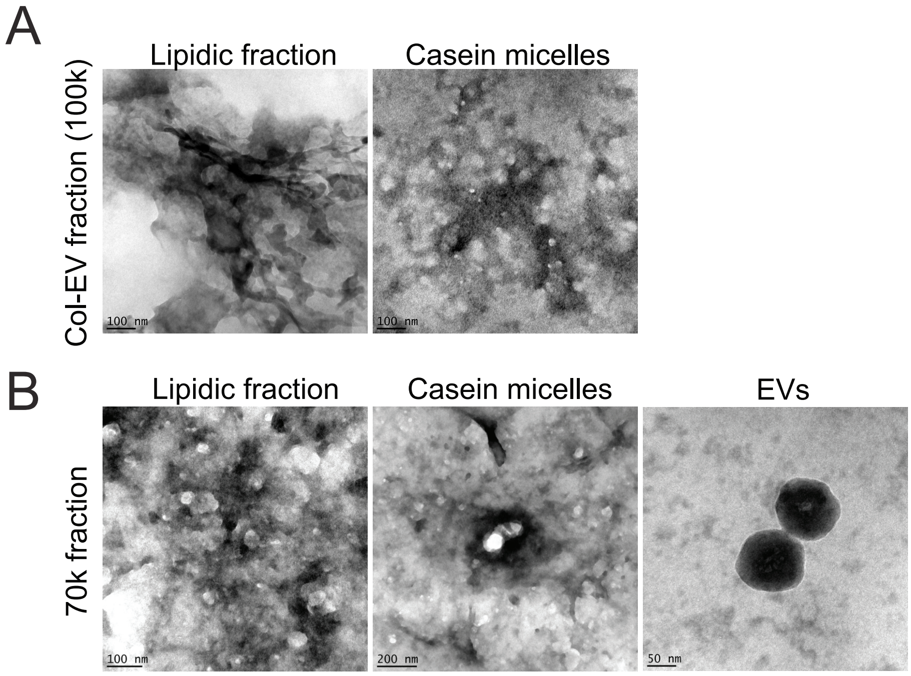


**Supplementary Figure 2.** Representative electron microscopy of different protein and lipidic fractions from ultracentrifugation products. **A)** Col-EV fraction (100k xg). **B)** 70k xg fraction.

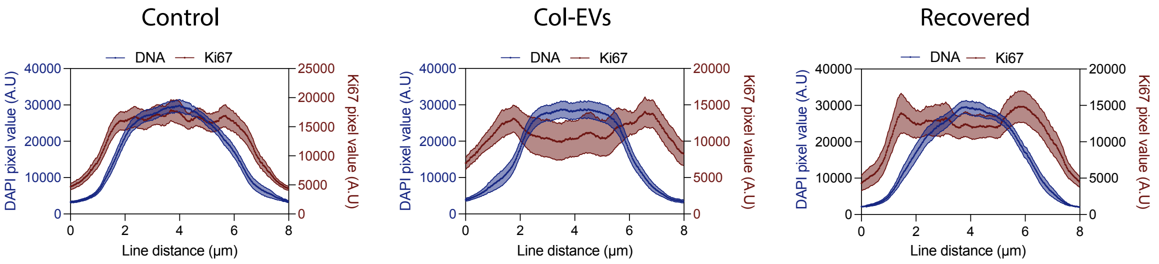


**Supplementary Figure 3.** Linescan measurements of metaphasic plates in LoVo cancer cells untreated, treated with Col-EVs or recovered after treatment. Blue lines indicate signal for DAPI linescans along metaphasic chromosomes. Red lines indicate signal for Ki67. Data indicates average +/- SEM, from N>20 positions/condition from n=2 independent experiments.

**Supplementary Movie 1.** Representative timelapse videos from LoVo and PANC1 control cells or treated with Col-EVs. Scale bar: 100 μm.
